# UPMaBoSS: a novel framework for dynamic cell population modeling

**DOI:** 10.1101/2020.05.31.126094

**Authors:** Gautier Stoll, Aurélien Naldi, Vincent Noël, Eric Viara, Emmanuel Barillot, Guido Kroemer, Denis Thieffry, Laurence Calzone

## Abstract

One of the aims of mathematical modeling is to understand and simulate the effects of biological perturbations and suggest ways to intervene and reestablish proper cell functioning. However, it remains a challenge, especially when considering the dynamics at the level of a cell population, with cells dying, dividing and interacting. Here, we introduce a novel framework for the dynamical modelling of cell populations packaged into a dedicated tool, UPMaBoSS. We rely on the preexisting tool MaBoSS, which enables probabilistic simulations of cellular networks, and add a novel layer to account for cell interactions and population dynamics. We illustrate our methodology by means of a case study dealing with TNF-induced cell death. Interestingly, the simulation of cell population dynamics with UPMaBoSS reveals a mechanism of resistance triggered by TNF treatment. This appoach can be applied to diverse models of cellular networks, for example to study the impact of ligand release or drug treatments on cell fate decisions, such as commitment to proliferation, differentiation, apoptosis, etc. Relatively easy to encode, UPMaBoSS simulations require only moderate computational power and execution time.

To ease the reproduction of simulations, we provide several Jupyter notebooks that can be accessed within a new release of the CoLoMoTo Docker image, which contains all required software and the example models.

## Introduction

Signaling pathways are organized in complex networks encompassing numerous cross-talks and feed-backs. Hence, the deregulation of one specific pathway often leads to non-intuitive effects. Mathematical modeling of such complex and intricate networks can help to understand and predict experimental results [1, 2, 3, 4, 5]. The choice of the mathematical formalism is made on the basis of the biological question, and the available data [6], and the nodes contained in the network correspond to genes or proteins whose activity shows an impact on the altered pathways and the biological responses [1].

It is well known that the micro-environment plays a key role in many pathologies, including cancer [7] and auto-immune diseases [8]. Hence, it is important to explore the dynamics of each cell type taking into account their micro-environment: how each cell type replicates, dies, and interacts with other cells. This requires a proper computational description of heterogeneous interacting cell populations.

To date, most existing mathematical models focus on the description of signaling networks at the level of individual cells. These networks are often depicted as wiring diagrams (or graphs), which describe the interactions and influences between genes and proteins inside the cells or in response to external cues (inputs of the network). However, the fate of individual cells will impact the fate of the population. It is thus important to consider cells dying, cells dividing and cells interacting or differentiating to fully understand the population dynamics.

Some modeling approaches have tackled this issue using an agent-based formalism: each agent corresponds to a cell, whose activity depends on that of its neighbors [9, 10] (i.e., CellSys [11], PhysiCell [12]). Attempts to explicitly include regulatory networks into these agents [13] have shown that this can easily become computationally costly or would increase considerably the number of parameters to tune (e.g., PhysiBoSS [14]). Other applications rely on cellular automata, e.g., Epilog [15], or yet on the Potts model, which can be also considered as an agent-based approach [16]. Finally, some works (e.g., BooleanNet [17], or [18]) apply stochastic and probabilistic simulations but on non-interacting populations of cells.

All these different methodologies have their own applications, e.g. CellSys, PhysiCell and PhysiBoSS for tumor growth, Epilog for spatial patterning, Potts models for vascularisation and angiogenesis [19], and are very promising to explore the effects of perturbations at the level of individual cells and understand how these perturbations impact cell populations. They constitute a first step towards the use of multi-scale modeling for clinical applications [20, 21].

Here, we present a modeling framework that consists of gathering all pathways in a single and unique model, even if they belong to different cell types, using a Boolean formalism. Throughout the article, we will refer to this idea with the term “MetaCell dynamics”, which cannot be considered as an agent-based model. The software tool UPMaBoSS (Update Population MaBoSS) was created to simulate the population dynamics of different interacting cell types keeping track of the individual gene/protein activity. UPMaBoSS uses MaBoSS grammar as a basis for the update of the Boolean values [22, 23].

The idea behind the “MetaCell dynamics” is to consider the whole micro-environment as a unique “cell”, or a meta-cell, which includes all the different cell types present in the population. It relies on a qualitative model of intra-cellular signaling networks built in Ma-BoSS (more details below), which enables probabilistic simulations of cellular models. UPMaBoSS alternates the simulation of intra-cellular models with regular updates of estimates of cell population size and environmental signals. These updates are based on the values of key network nodes (e.g., receptors and ligands) and processes accounting for cell division, cell death, and cell-cell interactions. The MetaCell dynamics can be considered as an heterogeneous cell population, withdifferent cell types, or cells in different states. The extension of pathway model(s) constructed in MaBoSS is easy, and the computational cost is only slightly larger than for MaBoSS original model(s). UPMaBoSS enables the description of complex cellular networks encompassing relatively large numbers of components (up to a few hundreds).

UPMaBoSS can be used for any biological questions that involve interactions between cell types. We anticipate that the framework could be applicable to new technologies such as single cell or CRISPR/Cas9 data. For single cell data, the parameters could be adjusted to the proportions of some cell types observed in different experimental conditions, or the increasing or decreasing probabilities of proliferating or dying cells could be verified for the CRISPR/Cas9 results.

We illustrate here our modeling scheme with a toy model and a case study dealing with the effect of TNF (Tumor Necrosis Factor) on cell fate decision following death receptor engagement.

To allow reproducibility of the results, we provide several notebooks as supplemental materials, with examples of models in MaBoSS language or in standard format such as SBML-Qual [24]. For the latter, the models need to be adapted to account for cell death and division, as well as for cell-cell interactions.

## Materials and methods

### Boolean logic

A network describing the interaction within an individual cell can be defined as an activity flow in SBGN standard (Systems Biology Graphical Notation [25]) and can be referred to as a regulatory graph or an influence network. In such networks, nodes represent molecular entities (genes, proteins, complexes) or processes, while edges denote the influence of source nodes onto target nodes. Any component of this cellular network can further influence the micro-environment by releasing signals (output nodes), while the microenvironment is sensed via receptors (input nodes). Our modeling scheme is based on Boolean networks which consist in discrete sets of Boolean variables: each variable (corresponding to the node of the network) has a *Boolean state*; a *network state* is, by definition, a Boolean vector representing the Boolean state of each node.

### MaBoSS description

Our modeling scheme relies on the previous development of MaBoSS [22, 23]. MaBoSS is a C++ software enabling the simulation of continuous/discrete time Markov processes, applied on a Boolean network. More precisely, a MaBoSS model is a set dynamical rules defining the requirements and the rates of each target node transition depending on the status of the source nodes. MaBoSS algorithm considers *network states*, i.e., Boolean vectors representing the Boolean state of each node, and applies a probabilistic scheme to this set of network states. In addition to the set of dynamical rules, a MaBoSS simulation requires an initial condition (an initial probability distribution over network states) and constructs a continuous time Markov chain. In practice, from an initial probability distribution, MaBoSS estimates the time-dependent probabilities of network states. These trajectories can be interpreted as a non-interacting desynchronized homogeneous cell population of a unique cell type, which entails important limitations when trying to describe the tumor micro-environment and the response to treatments. In practice, running MaBoSS requires two text files: 1) a file containing the description of the Boolean model of the cellular network to simulate (with a *bnd* extension, for *b*oolean *n*etwork *d*escription); 2) a file containing the parameters of the simulation (with a *cfg* extension, for *c*on*f*i*g*uration), including the specification of the initial state.

MaBoSS generates text files recapitulating simulation results, such as a list of fixed points, trajectories, time evolution of model state probabilities and stationary distributions (for more details, we refer the reader to the reference card of MaBoSS available at https://maboss.curie.fr).

### UPMaBoSS

In a simulation of a MaBoSS model, cells do not communicate, and at the end of the simulation, the cells can reach a “division” state (e.g., with a marker for proliferation ON) or a “dying” state (e.g., with a marker for apoptosis ON). To implement cell division and death explicitly in the simulations, we built UPMaBoSS that allows interactions between cells through the environment (with ligand-receptor interactions), and that can remove cells that are dying and double cells that are dividing. To encode this information, a new text file (with extension *upp* needs to be created and the MaBoSS files (*bnd* and *cfg*) need to be modified accordingly.

New nodes accounting for division and death are included in the MaBoSS *bnd* file and declared in the *upp* file. The status of the receptors depends on that of the environment. The receptors, initially considered as inputs of the MaBoSS model, inform on the microenvironment status and are updated according to MaBoSS external variables, whose update rule is given in the *upp* file and its initial value is given in the *cfg* file.

In MaBoSS, an external variable is, by definition, a variable that can be controlled externally and whose value is defined in *cfg* file. It is represented by a name starting with a $, and can be used anywhere in a transition rate formula (corresponding to the condition allowing the update of a node). In practice, receptor activation and/or inhibition rule(s) contain MaBoSS external variables defined in the *cfg* file; these external variables are updated by a specific formula that contains ligand probabilities, specified in the *upp* file. To summarize, two new nodes are added to account for death and division of cells, and the receptors are dynamically defined and are dependent on the availability of ligands in the environment. The initial condition, i.e., the initial probability distribution over the network states, is set up in the *cfg* file.

The population size is initially set to the arbitrary value 1. A MaBoSS simulation is run for a time length defined by the parameter *max time* in the *cfg* file. At the end of the first run, network state probabilities are updated: a) if the network state has the active death node, its probability is set to 0; b) if the network state has the active division node, its probability is doubled. These updated probabilities are re-scaled so that the sum of all network state probabilities remains equal to 1, and the population size is updated accordingly. These updated and re-scaled probabilities define a new probability distribution, used to update the external variables accounting for ligand-receptor interactions. This new probability distribution is injected as the initial condition for the next run (Figure 1). The process is repeated *n* times, where *n* is the number of steps defined in the *upp* file. A detailed description of the algorithm is provided in the supplementary material.

**Figure 1.**
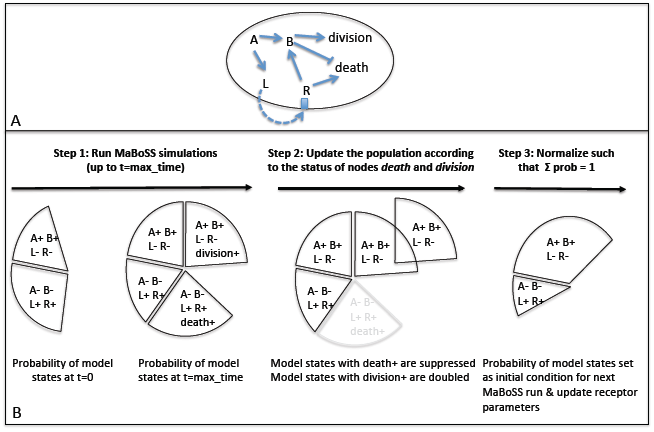
UPMaBoSS simulation. (A) An example of the MetaCell dynamics approach: A can activate B and release L (ligand) outside the cell, which activates R (receptor) on the membrane of the same cell; R can, on one hand, activate signaling pathways leading to cell death, and on the other hand, trigger pathways leading to cell division and inhibition of death through the entity B. (B) For the model shown in (A), UPMaBoSS algorithm applies several times the three following consecutive steps. *Step 1*: MaBoSS is run for a given initial condition and time length (only some selected model states are shown here: the state [A+,B+,L-,R-] means that only A and B are at 1 and L and R are at 0, the size of the pie portion corresponds to its probability); for some states, this leads to the activation of the nodes division or death. *Step 2*: After the MaBoSS run, state probabilities are updated by removing those that have the death node active and doubling those that have the division node active; the size of the population is multiplied by the sum of these new probabilities, which are then normalized. *Step 3*: The transition rates of the receptors are updated with the newly computed probabilities; the next MaBoSS run starts with the normalized probabilities as initial condition. The number of runs *n* is defined by the user and this sequence of three steps is thus repeated *n* times.

### Usage

UPMaBoSS is an executable file than can be run in a unix environment in which MaBoSS is accessible. A Python package (integrated in “pymaboss”) has been developed and is available within the CoLoMoTo interactive notebook. [26, 27]. In practice, there are four ways to simulate a population model dynamics using UPMaBoSS: (1) using a Docker image, which necessitates to install Docker locally; (2) creating a conda environment and then using a jupyter notebook or launching UPMaBoSS directly with a command line; (3) running the notebooks directly in the browser via binder (instructions are in the GitHub of UPMaBoSS, see below); and (4) using MaBoSS 2.0 environment (UPMaBoSS belongs to the tools in MaBoSS package that can be downloaded at https://maboss.curie.fr/UPMaBoSS). All details and commands are provided in the supplementary material and in the GitHub repository: https://github.com/sysbio-curie/UPMaBoSS-docker.

## Results

### Definition of a *MetaCell*

A model is a set of variables (one for each of the nodes of the regulatory network describing the interactions between genes, proteins or complexes) and the value of each of these variables is controlled by a logical rule.

In the MetaCell (a model “beyond” a simple cell model), as we define it, we consider that all signaling pathways of interest are gathered in a unique cell model, thus possibly mixing different cell types, which considers that, all pathway components are potentially present in each cell. The MetaCell idea consists in representing different cell states as different configurations of values of these components. The biological focus of a model of the MetaCell is to describe the interactions between signaling pathways that lead to the activation of markers of proliferation (e.g., transcription factors of the cell cycle as E2F1, E2F3 or the cyclin/CDK complexes), apoptosis (e.g., p53 or the caspases), or other phenotypes such as senescence (e.g., p16), angiogenesis (e.g., VEGF). In particular, MetaCell modeling can reveal how perturbations of a normal condition may affect the activity of these markers, knowing that the activation of the signaling pathways inside a cell may depend on the micro-environment and the status of its neighbors.

The advantage of this framework is that it can model different interpretations of the impact of the microenvironment on individual cell fates and on the population dynamics. This can be exemplified by the two following different points of view. The first one hypothesizes that all cell types are already known, that the cells have already differentiated. In this case, we simply merge the cell types into a model of a MetaCell (Figure 2, A). The second point of view is that we can consider undifferentiated cells with all possible signaling pathways inside each cell. As the cells differentiate, some will activate some pathways, other will activate other pathways (Figure 2, B).

**Figure 2.**
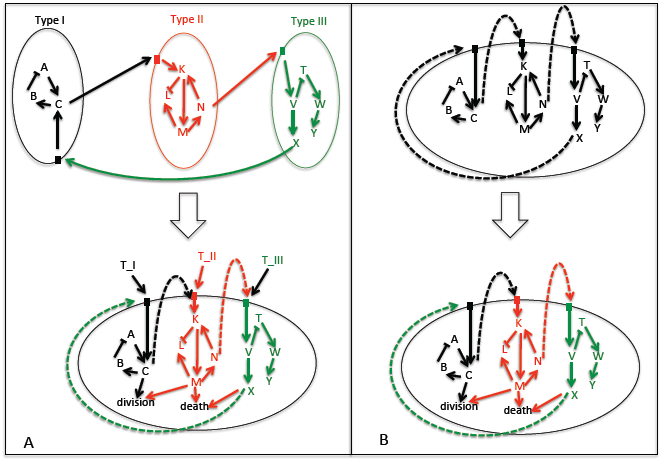
The MetaCell. A network of a MetaCell is constructed by regrouping all the signaling pathways that can be activated in different cell types, including ligand-receptor interactions. Two cases are considered: (A) when the cell types are already known and integrated into a single network, and (B) when the cell is initially undifferentiated and gets differentiated into different cell types (note that it is not explicitly shown in the figure but one given pathway could be activated into two different cell types, see text). In both cases, cells can die, divide or interact through ligand-receptor interactions.

To allow some flexibility in the simulation of different cell conditions, we suggest to add a node that represent the cell type and that will allow the components of this cell type to be active or not in a simulation. For the first point of view (Figure 2, A), these nodes are static variables, for the second point of view (Figure 2, B), they are dynamical variables. For instance, if in a MetaCell, T cells and tumor cells are included, we can add two nodes *T cell* and *Tumor* linked to all components of the pathways of the corresponding cell type. If one simulation represents the fate of a cell type, both cell types cannot be initially set to 1 at the same time. The population will be a mixture of T cells and of tumor cells. But, both cell types will be able to communicate, die or divide. Of course, a MetaCell can mix these two points of view: some cell types may change over time and other may not.

One diffi culty may arise when pathways that are gathered inside a MetaCell have common parts. This situation could not be solved by an automatic algorithm, because biological knowledge should be used for fusing pathways: if two pathways have common nodes that are, in fact, different entities, these common nodes need to be renamed; if the common nodes are indeed the same entities, dynamical rules that update these nodes should be modified accordingly. See below (“The dynamics of a MetaCell in UPMaBoSS”) for some hints.

### The dynamics of a MetaCell in UPMaBoSS

The UPMaBoSS (for Update Population MaBoSS) framework enables the exploration of the dynamics of a MetaCell and produces time-dependent probabilities of states for each cell type, together with time-dependent population size, allowing an interpretation of the dynamics at the level of the cell population, considering that:

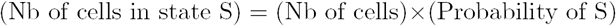

UPMaBoSS is a modified and extended version of MaBoSS software [22, 23]. In the model definition, two nodes are added: one node accounting for cell division, and another one for cell death. The pathways in the MetaCell are described in MaBoSS framework according to the following rules: each node of the network represents a gene, protein, complex, phenotype or cell type; the nodes can only have one Boolean value; every MetaCell state is represented by a set of Boolean values associated with the set of nodes (including the division node, the death node, the ligand and the receptors, cell type nodes); the logical connections between nodes are described by two transition rates (activation/inhibition) associated with every node, formulated in the language of MaBoSS; for autocrine/paracrine loops, the rates associated with the update of the state of the receptors should contain term(s) that depend on the MetaCell state probabilities. The setting of initial conditions is the same as for MaBoSS. As described above, all pathways are gathered in a single model.

If pathways share common nodes, biological knowledge should be used to do this fusion correctly: a) if the common nodes are different entities, they should be renamed properly; b) if the common nodes are activated/inhibited in different situation corresponding to different differentiated cell types, the respective rates of transitions should be linked to the respective cell type node and fused into a single rule (*eg rate up* = (*NodeTypeA*?*V alueAInTypeA* : 0) + (*NodeTypeB*?*V alueIntypeB* : 0)); c) if the common nodes do not belong to the two cases above (e.g., cell types in the process of differentiation), the rates of transition should be combined in a more complicated manner, based on the biological knowledge, using the flexibility of MaBoSS language for transition rates.

The outputs obtained with UPMaBoSS include time-dependent relative population sizes (that are compared to the population size at t = 0), MetaCell state probabilities including the status of death, division, cell types (Figure 3).

**Figure 3.**
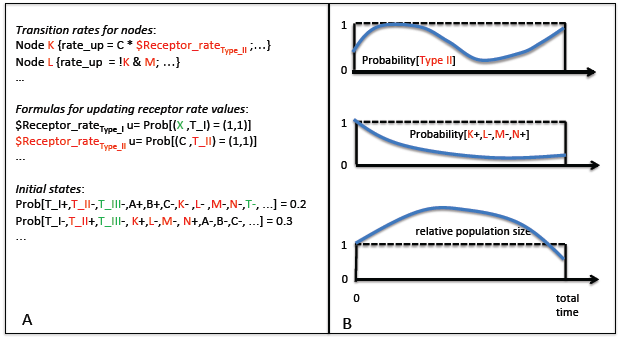
Inputs and Outputs for UPMaBoSS. (A) Inputs needed for a simulation with UPMaBoSS: transition rates for nodes of the MetaCell, Formulas for updating receptors rates values, and initial conditions. The colors correspond to the cell types presented in Figure 2. (B) Ouputs of UPMaBoSS: time-dependent probabilities of cell types (example of cell type II from Figure 2, and of the corresponding model states, and time-dependent relative population size.

UPMaBoSS launches several consecutive MaBoSS runs (defined by the user). At the end of each run, the population is updated synchronously: new model states are produced according to the parameters influencing the population status (death, division, receptor activity), setting a new initial condition for the next MaBoSS run (see Material and Methods).

To properly simulate the population dynamics, there are important parameters to define, which can be separated into two families: the ones that have a biological interpretation and the ones that are used for the simulations.

The first family of parameters include the rates of activation or inactivation of a variable, which can be derived from experiments. They can correspond to the time necessary to achieve transcription, (de)phosphorylation, synthesis or degradation, when the information is available. The transition rates can also be separated into fast or slow variables. If the information is not known, the default value 1.0 is used. Other parameters can account for the initial conditions or the length of the experiment (total time of simulation is reached when (total time) = (number of steps) × (length of MaBoSS run)).

The second family of parameters informs on the number of trajectories to include in the computation, or the length of one MaBoSS run (to ensure, in some cases, that transitory behaviors are observable, see an example below).

An exhaustive list of these parameters is provided in supplementary material, including default values and hints for choosing the values of these parameters. In some cases, a sensitivity analysis might be helpful to search for the appropriate range of parameter values. An example of varying the length of one MaBoSS run *max time*) and its impact on the expected results is studied in a dedicated python notebook (supplementary material).

### Case studies

We illustrate the use of UPMaBoSS with two examples, the first one is a toy model of cell-cell interactions, and the second one is based on a published model of cell fate decision process in response to death receptor activation [28].

#### Toy model

As a first application, we present a toy model to highlight that different dynamics can be obtained if either a unique cell or a MetaCell are considered. The model illustrates a differentiation process into two cell types, T1 and T2. The differentiation is initiated by a trigger T. A part of the signal goes through a ligand L that activates the receptor R. A single model to simulate both cases is defined and shows the results of the simulation first when considering a unique cell (or a population of independent cells), and then taking into account the status of the population with cell-cell interaction (ligand to receptor). The MaBoSS external variable $*innerOn* is used to distinguish between the two cases: in the case of the unique cell when $*innerOn* = 1, R can be activated by L directly inside the cell; in the case of the MetaCell, when $*innerOn* = 0, R activation rate is proportional to the probability of active L, through the update function of $*outerL* described in the *upp* file (Figure 4A).

**Figure 4.**
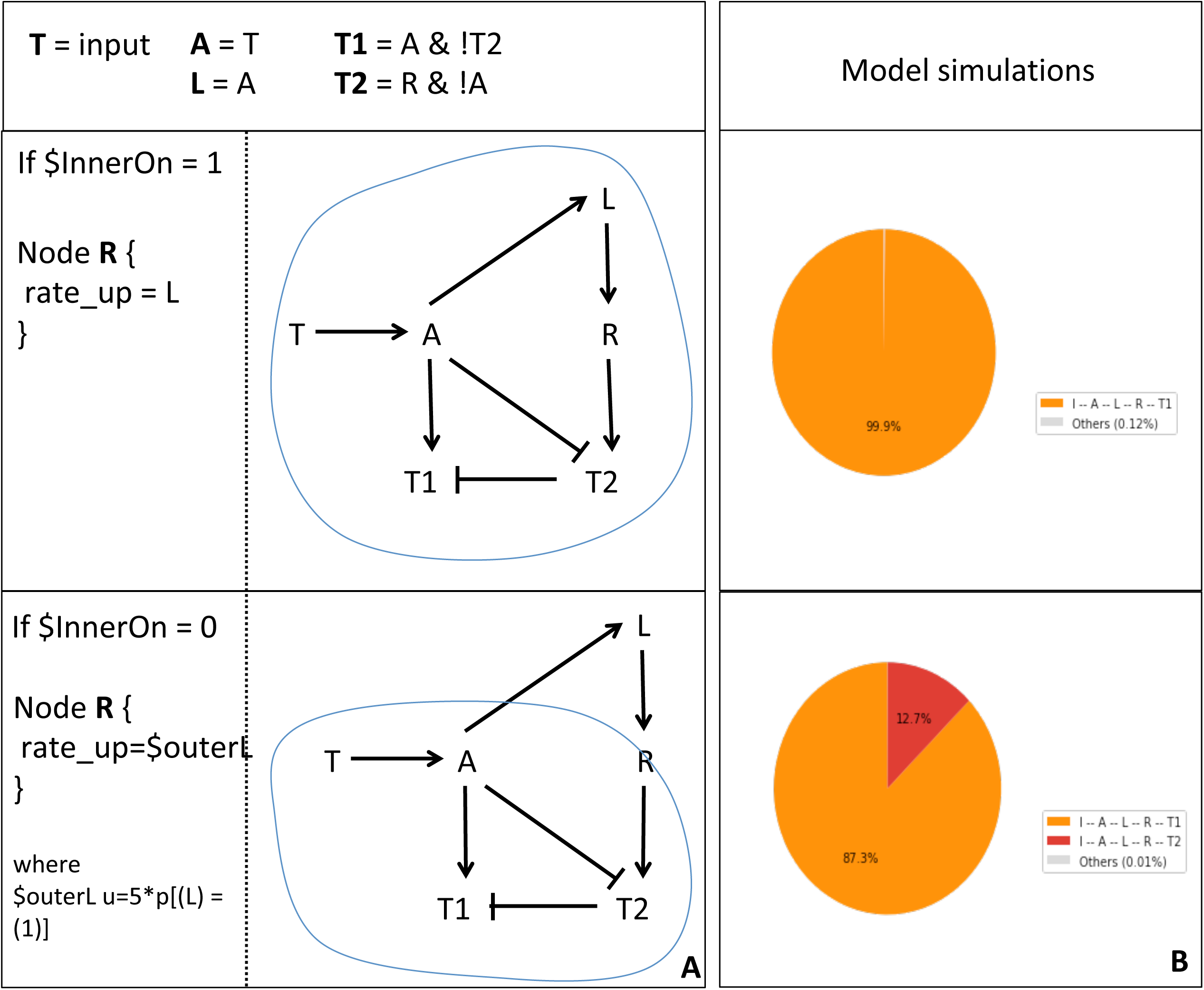
Toy model. (A) Definition of the toy model with logical rules (upper panel) and conditional rule for R depending on the value of the external parameter innerOn. If innerOn is equal to 1, then A is able to activate L in all cells (middle panel). If innerOn is set to 0, then the value of R will depend on the population status of L (lower panel). (B) Model simulations of the two cases: when innerOn=1, only T1 cell type can be reached; when innerOn=0, a proportion of cells can differentiate into T2 cell type.

When the unique cell is considered, T2 cell phenotypes can never be reached because A is always present when R is active and thus, inhibits T2 (Figure 4B, upper panel). When the population model is considered, R is updated according to the population state of L; consequently, R can be activated in some cells that do not have active A, producing the differentiation in T2 cell type with a non-zero probability (Figure 4B, lower panel).

A jupyter notebook is provided for this example and shows not only the UPMaBoSS model with the three files but also a way to build a population model from a standard model format (here a bnet format[29]).

#### TNF treatment

This case study illustrates how a cell population responds to different protocols of a drug treatment. For that, the model needs to encompass pathways regarding cell proliferation and death, because these phenotypes act directly on the cell population size.

#### Description of the cell fate decision model

We focus here on a model initially built to understand how cell death receptors engagement can lead to three different cell phenotypes: survival with the activation of NF*κ*B pathway, necrosis, and apoptosis with the cleavage of caspase 3 [28]. The original analysis explored which components contribute to each phenotype, as well as the interplay between the three pathways. In particular, the three pathways exhibit mutual inhibitions, thereby ensuring that the corresponding cell fates occur exclusively from each other. A slightly modified version of the model is available on the MaBoSS web page (https://maboss.curie.fr) and on the GINsim repository (http://ginsim.org/sites/default/files/CellFate_multiscale.zginml), with the parameter choices described in the supplementary material.

In the present study, we aim at completing this analysis by considering the impact of the timing and duration of TNF treatments. For this analysis, the model was extended by adding a feedback from NF*κ*B pathway to TNF*α* (Figure 5, A). Indeed, it has been showed that TNF*α* is a target of NF*κ*B, and that constitutive activation of NF*κ*B leads to systemic inflammation through TNF*α* activation ([30], and in primary macrophages [31, 32, 33]). We further decided to focus on the role of TNF*α* and thus kept Fas OFF for all our simulations.

**Figure 5.**
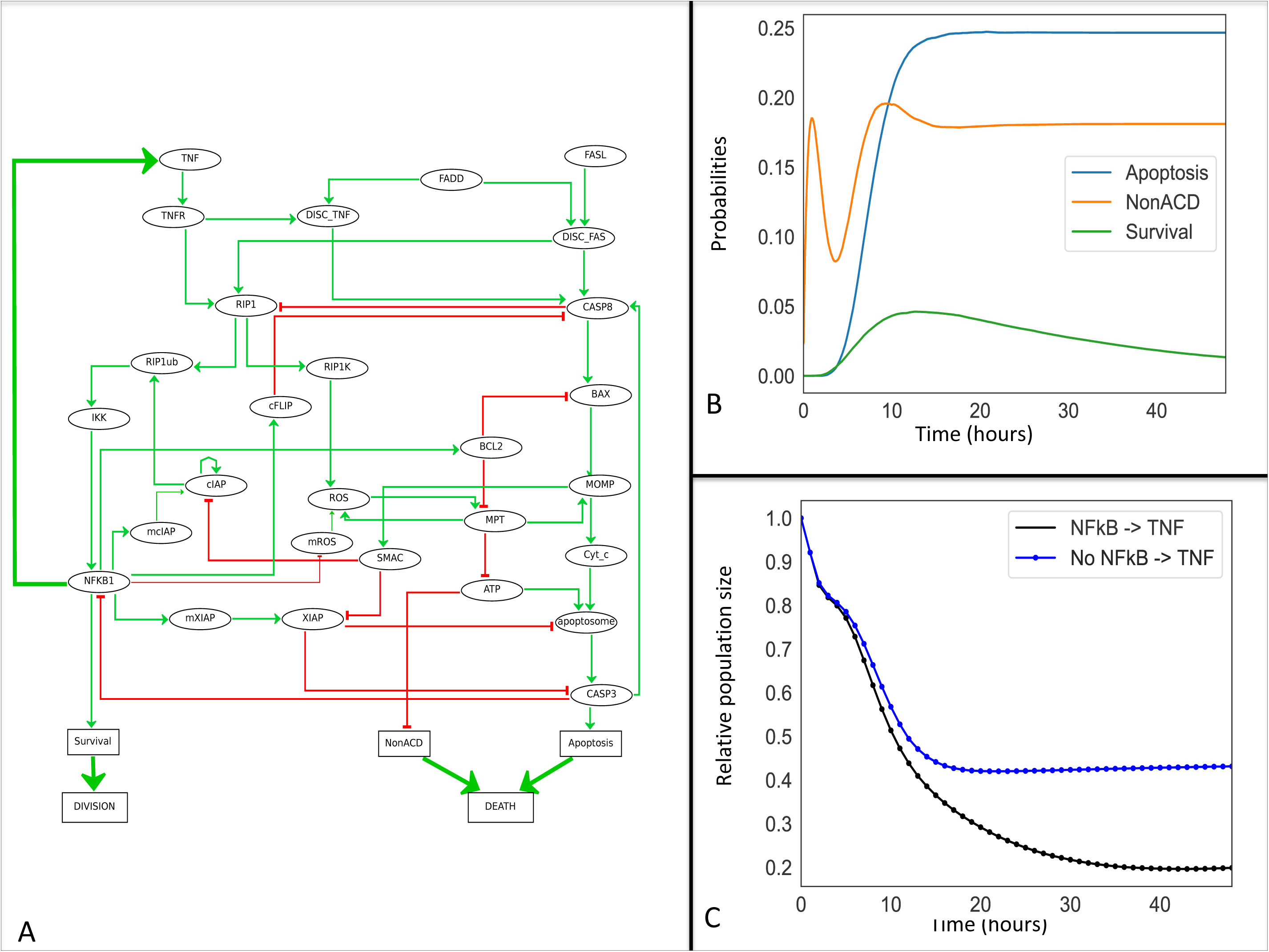
Cell fate model for TNF*α* resistance. (A) The model has been slightly modified from [28]. Some nodes representing the mRNA of cIAP, ROS and XIAP family members have been added. The circle nodes represent genes, mRNA, proteins, or complexes, and rectangular nodes account for the activity of phenotypes. Green and red arrows represent positive and negative influences, respectively. The thick green arrows denote activating interactions added to the initial model: a feedback from NF*κ*B to TNF*α* encodes the ligand-receptor activation, while the “Division” and “Death” nodes have been introduced specifically for UPMaBoSS population updates. (B) Simulation of the cell fate model with MaBoSS for a length equivalent to 48 hours. (C) Simulation of the model with UPMaBoSS: relative population size with (black) and without (blue) the TNF paracrine signaling.

We explored the effect of sequential treatments of TNF*α* at a cell population level. Interestingly, several studies showed that prolonged treatments of low doses of TNF lead to resistance in prostate patients [34], and that TNF exhibits a dual role in tumor progression: at low doses, it triggers angiogenesis [35], whereas at high doses, it induces cell death, mainly through necrotic effects [36].

#### Biological questions

We simulated the model for a period corresponding to 48h of cell culture (unit time set to one hour). Experimentally, there is no consensus for time duration of TNF effect *in vitro*; nevertheless, some effects need more that 24 hours to appear [37], which justifies the choice of 48 hours. This is confirmed by the simulations within MaBoSS and UPMaBoSS below (Figure 5) because stability is reached after 48 hours. For transient activation of TNF, we considered a TNF half-life of 4 hours (degradation rate of 1*/*4). As TNF degradation rate varies extensively depending on experimental conditions, we select four hours as a reasonably small interval within 48h.

This simulation is particularly important to define the time step for each MaBoSS run when using UPMaBoSS. Indeed, the chosen time window (i.e., max time) must be such that the population is in a transient state. The MaBoSS run indicates that the best value for the parameter max time needs to be set to 1 hour (just before the peak of activation of (Figure 5, B). To simulate the population dynamics, we thus proceed to compute 48 MaBoSS runs with max time equal to 1 for each run. We explored the effect of choosing a different value in the supplementary material (jupyter notebook *TimeStepDependency*.*ipynb*).

During the MaBoSS simulation, we noticed that non-apoptotic cell death first decreases before increasing to reach a steady state solution after t = 15. This dynamics is due to the activity of ATP, itself dependent of that of RIP1. RIP1 increases until CASP8 is activated and able to inhibit it. It takes longer to activate CASP8 than RIP1. This behaviour is typical of incoherent feedforward loops [38].

For this biological application, we focused on two questions:

1. What is the effect of the feedback from NF*κ*B to TNF*α* at the population level when treated by a transient activation of TNF*α*?
2. What is the effect of TNF*α* sequential treatments on the population dynamics?

Two model variants were thus considered: with and without the NF*kappa*B → TNF paracrine loop, with a transient TNF treatment (Figure 5, C). For the model with the paracrine loop, we further studied the following scenarios (Figure 6) where cells are treated with:

**Figure 6.**
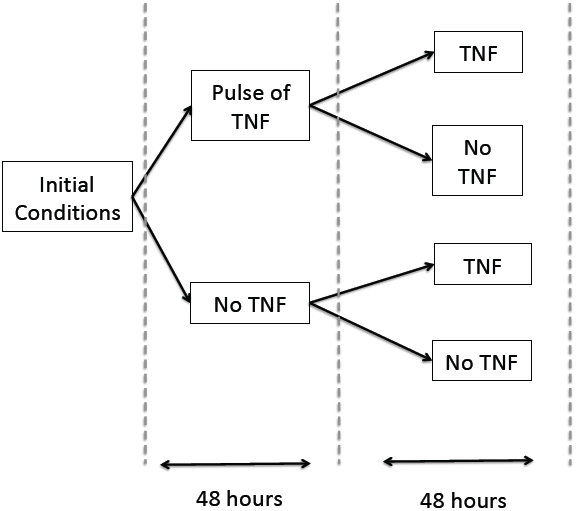
Simulation scenarios for the study of TNF resistance.

(a) a transient TNF treatment at time 0, followed by a constitutive TNF treatment at 48 hours (“Pulse TNF” + “TNF”); (b) no TNF treatment at time 0, followed by a constitutive TNF treatment at 48 hours (“No Pulse TNF” + “TNF”); (c) a transient TNF treatment at time 0, followed by no TNF treatment at 48 hours (“Pulse TNF” + “No TNF”); and (d) no TNF treatment at time 0, followed by no TNF treatment at 48 hours (“No Pulse TNF” + “No TNF”).

#### TNF treatments in wild type conditions

Simulations of the temporal evolution of cell populations are displayed in Figures 5(C) in presence or absence of the feedback and in Figure 7 for two different TNF treatment scenarios.

**Figure 7.**
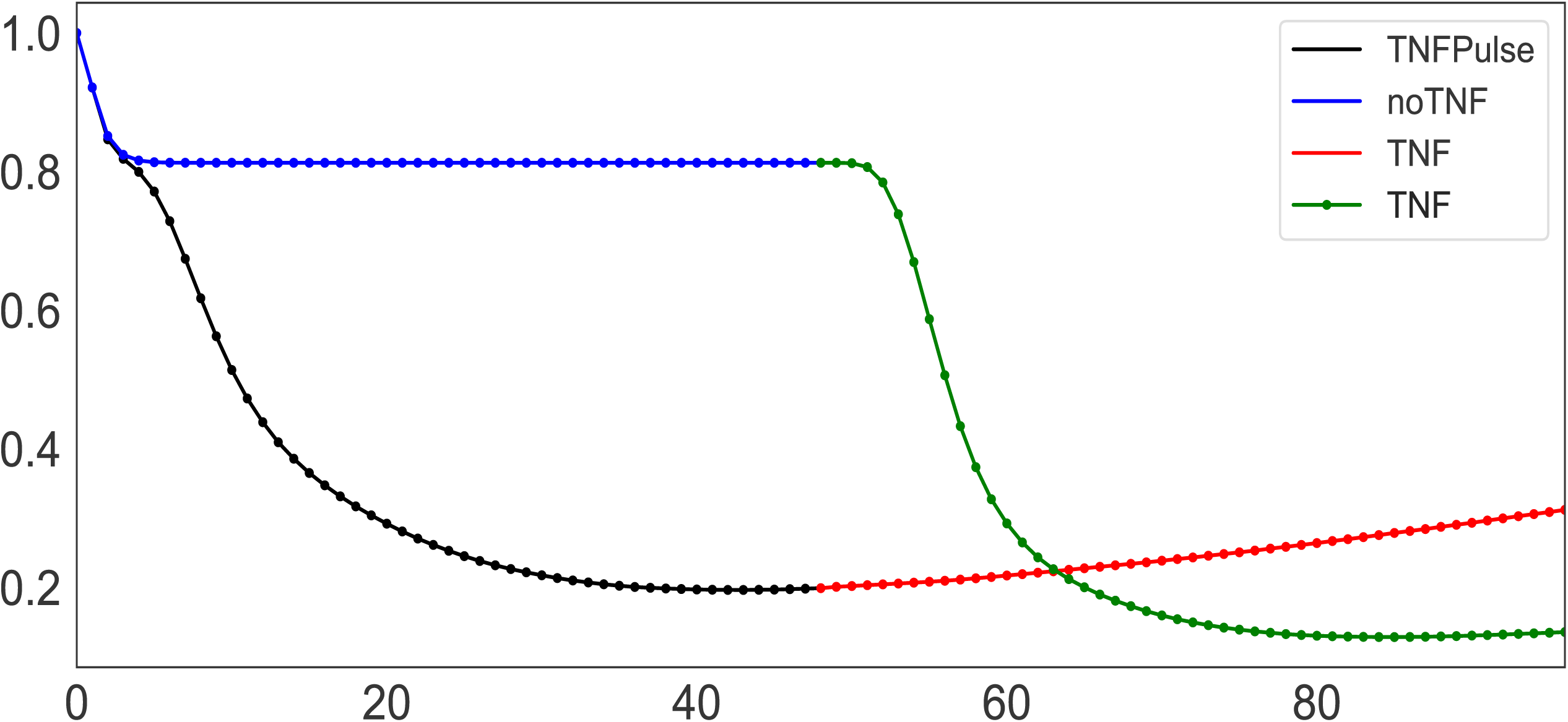
Growth curves for different TNF treatment scenarios. One scenario corresponds to the simulation of cells initially treated by a pulse of TNF (black segment), followed by a constitutive TNF treatment at t = 48 hours (red segment). The other scenario corresponds to the simulation of cells initially untreated (blue segment), but receiving a constitutive TNF treatment at t = 48 hours (green segment).

In Figure 5(C), following a pulse of TNF at t = 0, the comparison of population growth curves in the absence (blue curve) or in the presence (black curve) of the feedback from NF*κ*B to TNF*α* indicates that the TNF paracrine loop leads to a decrease of the population size (from 0.43% to 0.20% of the initial population size of 100%).

Remarkably, when a sustained treatment is applied, the impact on the population size differs depending on whether the population has already been treated or not, in a non-intuitive way. Indeed, as shown in Figure 7, after 48 hours, the population initially untreated (referred to as “noTNF” + “TNF”, blue + green curves) decreases faster than the cell population initially treated with a pulse of TNF (referred to as “TNFPulse” + “TNF”, black + red curves). This difference could be interpreted as a resistance mechanism: cells that have already been exposed to TNF, even transiently, do not respond as well to a second TNF treatment compared to cells that have never been treated with TNF. Noteworthy, this “resistance” results purely from network dynamics, in the absence of any genetic modifications.

#### TNF treatments in mutant conditions

The analysis was done on the wild type conditions, but in cancers, the many different mutations found in patients may affect the efficacy of the response to some treatments. To explore the potential roles of the different model components in the observed TNF resistance, we simulated the effects of all possible single mutants, following the *in silico* protocol illustrated in Figure 6. For each single mutant, we measured the population ratio at t = 96 hours for the four scenarios: “No TNF” + “TNF”, “Pulse of TNF” + “No TNF”, “No TNF” + “No TNF”, and “Pulse of TNF” + “TNF”.

All single mutants were computed (Jupyter note-book *CellFateModel_upmaboss* in supplementary material). We illustrate here the results for three genetic backgrounds: wild type, IKK knock-down, and RIP1K knock-down (Figure 8) with the four protocols (Figure 6 focusing on the period between t = 48 to t = 96. We only focus on the response after t = 48 hours for simplicity, and to improve the comparison, we normalized the population ratio found in Figure 7. Note that two conditions were added to those shown on Figure 7, where only two of the four protocols were simulated, i.e., we added the cases when there is no treatment after 48 hours, no matter what the cells receive at t = 0 (dashed lines).

**Figure 8.**
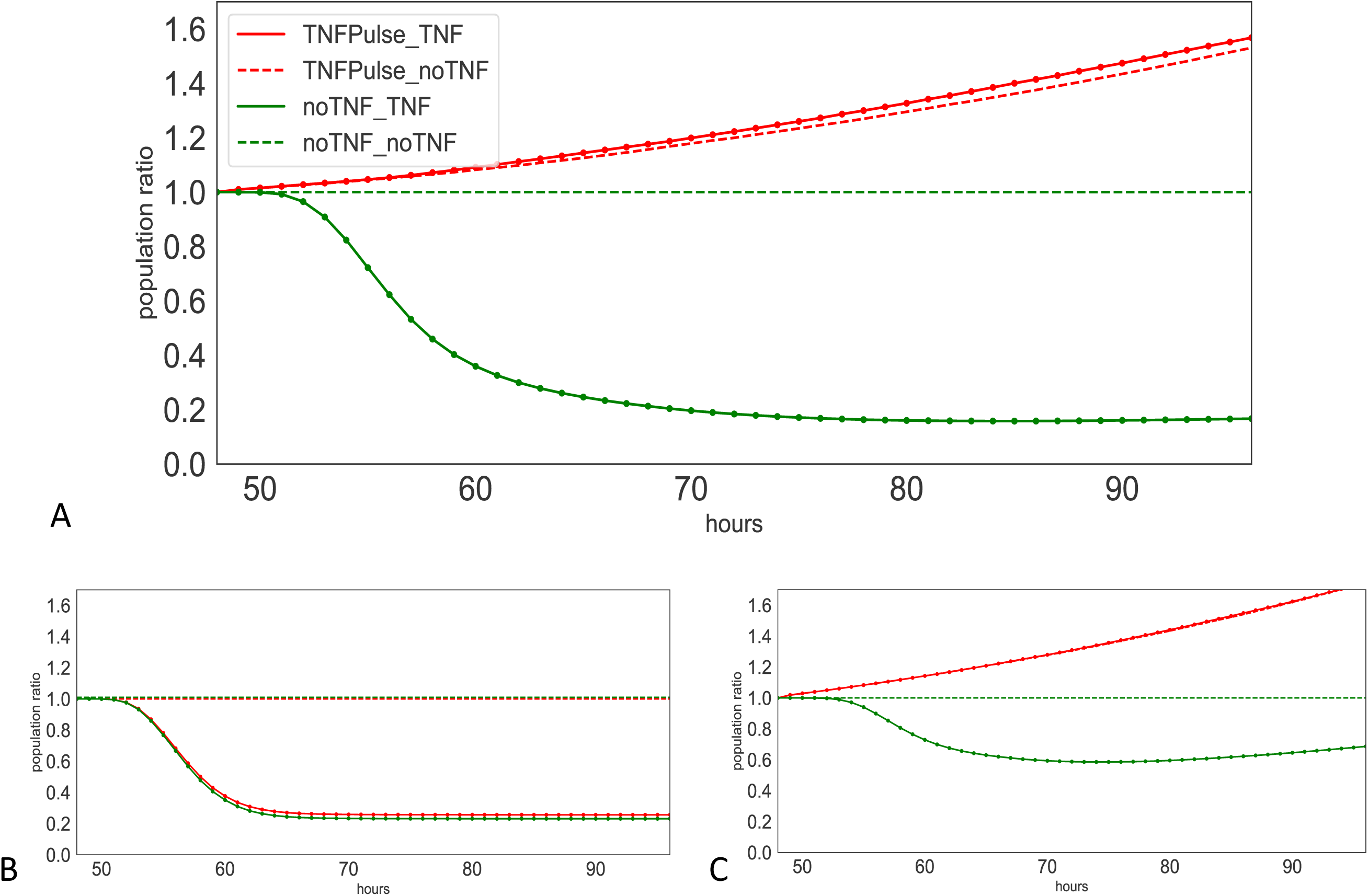
Population ratio at from t = 48 to 96 hours for three models. Population ratios for the four conditions described in the *in silico* experiments (Figure 6 in (A) wild type, (B) IKK knock-down, and (C) RIP1K knock-down

The wild type model clearly exhibits a resistance: when the cells have received a treatment, they do not respond whether they receive a second treatment or not (red curves representing increasing population ratio), whereas the plain green curve decreases (Figure 8 A). Note that the dashed green curve corresponds to the population of wild type cells without treatment (no signal) normalized to 1. The resistance effect observed in wild type is lost for IKK knock-down (Figure 8 B), as the two second treatments have no effect on the population ratios (plain and dashed lines coincide). For the RIP1K knock-down, the population reduction by TNF is not as strong as for wild type model, but the resistance mechanism is still observed (Figure 8 C).

IKK belongs to the NF*κ*B pathway that induces survival. This pathway participates in a positive feedback at the cell population level, which may explain why IKK knock-down shuts down the resistance to TNF by shutting down this feedback loop. RIP1K induces non-apoptotic cell death by blocking ATP, which explains RIP1K knock-down reduces the apoptotic effect of TNF (see *CellFateModel_upmaboss*).

Similarly, the effect of double mutations could also be studied. Note that the computation time is significantly longer when simulating the four protocols for all the double mutations. For our example, double mutant simulations do not result in additional insight because the single mutations are already informative, but it might be the case for more comprehensive models.

All results and figures of this analysis can be reproduced with the Jupyter notebook *CellFateModel_uppmaboss*.*ipynb*, which is provided as supplementary material, and can also be found in the folder *usecases* in the CoLo-MoTo Docker image, and or yet the address: https://maboss.curie.fr/UPMaBoSS/).

## Discussion

UPMaBoSS is a new a modeling framework enabling the exploration of cell population dynamics. It considers the division, the death and interactions of populations of cells. It is run using a stochastic approach but updated at regular intervals.

More specifically, a logical model is constructed from the fusion of signaling pathways, which may belong to different cell types, integrated into a single model description (referred to as MetaCell here) within Ma-BoSS language; cell-cell interactions, conditions for cell division and for cell death are added on top of the regular MaBoSS model.

Because UPMaBoSS ignores the spatial dimension, computational cost does not increase a lot compared to prior MaBoSS signaling models: because UPMaBoSS is made of consecutive MaBoSS runs, it is only the updating algorithm between each MaBoSS run that adds computational cost. We notice that the latter is small. Indeed, MaBoSS constructs more than ten thousands trajectories, as the updating algorithm just parses and writes probability distributions.

Using a simple toy model, we showed that the results of the simulations are different if we consider a homogeneous non-interacting cell population or if we consider that cells can communicate.

We applied our approach to a model of TNF induced cell fate. We show that the paracrine loop through NF*kappa*B enhances the death of cells through TNF effect. Surprisingly, this model shows a resistance mechanism to TNF treatment: once the cells have been treated by TNF transiently, they do not die upon a second treatment; and this effect is not due to (epi-)genetic selection.

UPMaBoSS framework produces results that can be validated experimentally by different techniques: the probabilities of the nodes corresponding to proteins/mRNA can be compared by experimental concentration (measured by western blot, immunofluoresence, qPCR, etc.). The probabilities of network states can be compared with the different techniques regarding cell populations: after labeling some proteins by fluorescent antibodies, flow cytometry or microscopic images can be used; the new single cell technologies could, in principle, be applied, by matching the population size to the different cell types.

We expect that our approach is suitable for modeling the effect of the micro-environment on cell fate for two reasons: an easy implementation on top of the construction of single cell type(s) model(s), and a low computational cost. Indeed, once pathways for individual cell types have been properly translated in MaBoSS language, they are gathered in a single model description. Then, the user needs to set up conditions for cell division and cell death, and for ligand-receptor interactions. The setting of the heterogeneous cell population description and proportion can be handled from the proper definition of initial conditions. For this case, we suggest to add nodes that represent cell types: when these nodes are initially set to 1, the cell type can be considered present, with the desired proportion.

We think that this approach is appropriate to model many processes in cell biology, including cell differentiation, innate/adaptive immune system activation, cancer micro-environment, tissue homeostasis, etc. The fact that the spatial dimension is not taken into account here might appear as a limiting factor in clinical applications. However, as a counter-example, some works have shown that the presence, more than the precise localization, of immune infiltrates in tumors constitute good prognostic biomarkers [39]. The important part lies in the translation of inter-cellular communication in terms of the ligand-receptor description used in UPMaBoSS. The implementation in docker or conda environment makes its use easy and flexible, as presented in the TNF treatment example within a Jupyter notebook.

One limitation of our approach is the somewhat artificial setting of population update time. However, the case study presented here suggests that results are moderately sensitive to changes of this parameter within a reasonable interval.

Another limitation is that UPMaBoSS is designed for signaling pathways, not for metabolic ones. However, the dynamics of metabolites could be added, if they play a role in the signaling pathways. For that, metabolites are represented by Boolean nodes, the level of metabolite(s) are therefore discretized and then “Booleanized” (transforming a multi-level node by multiple Boolean nodes) for MaBoSS format. The translation in MaBoSS will require some considerations about the timing and the dynamics of these metabolites, e.g., the consumption of reactants will be translated by an ultra fast degradation rate of a reactant induced by its products.

In summary, we provide a tool enabling the simulation an interacting heterogeneous population of cells allowing to take into account the influence of the micro-environment on the fate of a cell population. The quantitative output of UPMaBoSS facilitates the comparison to experimental observations and the consequent tuning of the parameters, which ultimately will provide finer model predictions.

## Author’s contributions

GS and EV conceived the algorithm and the tool. VN, EV and AN participated in the development of the tool. GS, AN, DT and LC worked on the examples. DT, EB, GK, GS, LS designed and supervised the study. GS and LC wrote the article but all co-authors contributed to the writing and agree on the content.

## Acknowledgements

We would like to thank I. Martins. GK is supported by the Ligue contre le Cancer (équipe labellisée); Agence National de la Recherche (ANR) – Projets blancs; ANR under the frame of E-Rare-2, the ERA-Net for Research on Rare Diseases; AMMICa US23/CNRS UMS3655; Association pour la recherche sur le cancer (ARC); Association “Le Cancer du Sein, Parlons-en!”; Cancéropôle Ile-de-France; Chancelerie des universités de Paris (Legs Poix), Fondation pour la Recherche Médicale (FRM); a donation by Elior; European Research Area Network on Cardiovascular Diseases (ERA-CVD, MINOTAUR); Gustave Roussy Odyssea, the European Union Horizon 2020 Project Oncobiome; Fondation Carrefour; Institut National du Cancer (INCa); Inserm (HTE); Institut Universitaire de France; LeDucq Foundation; the LabEx Immuno-Oncology (ANR-18-IDEX-0001); the RHU Torino Lumière; the Seerave Foundation; the SIRIC Stratified Oncology Cell DNA Repair and Tumor Immune Elimination (SOCRATE); and the SIRIC Cancer Research and Personalized Medicine (CARPEM).

## Additional Files

Additional file 1.

**Jupyter notebook** *ToyModelUP*.*ipynb*.

Additional file 2.

**Jupyter notebook** *CellFate upmaboss*.*ipynb*.

Additional file 3.

**Jupyter notebook** *TimeStepDependency*.*ipynb*.

