## Supplementary Material for "UPMaBoSS: a novel framework for dynamic cell population modeling"

#### 1 Description of UPMaBoSS

UPMaBoSS is a perl or python script that produces multiple runs of MaBoSS (or "steps"), in order to provide a modelling framework at population level. UPMaBoSS needs three files: a *bnd* file for the model description, a *cfg* file for the configuration file for each MaBoSS runs and an *upp* file that defines the different parameters describing the cell population. Note that all nodes must be set as external (in the *cfg* file).

##### 1.1 The *upp* file

This file needs to contain:

- A line defining the model node (present in the *bnd* file) that launches cell division, *e.g.*

  ```
  division = Division_Model_Node;
  ```

- A line defining the model node (present in the *bnd* file) that launches cell death, *e.g.*

  ```
  death = Death_Model_Node;
  ```

- A line defining MaBoSS executable name, *e.g.*

  ```
  MaBoSS = MaBoSS_custom_name;
  ```

- A line defining the number of steps, *e.g.*

  ```
  steps = 16;
  ```

This file can also contain line(s) defining external variable(s), used in the *bnd* file, that are updated at each step. Each of these lines should start with

the name of an external variable, followed by the update operator `u=` and an expression using the same operators than in MaBoSS language:
`+,-,*,/,AND,OR,XOR,NOT,(?:)`.
Probabilities of different states can be inserted, with following syntax:
`p[(node1,node2,...)=(1,0,...)]`.
Two keywords can be inserted: `#rand` representing a random number between 0 and 1 and `#pop_ratio` representing the population ratio. For instance:

```
$ExtVar1 u= 5*(p[(Node_A,Node_B,Node_C) = (0,1,0)])
```

Note that each of these external variables needs to be also defined in the *cfg* file, for the initial step of UPMaBoSS.

#### 1.2 Algorithm

For the step 0, UPMaBoSS runs MaBoSS according to the *bnd* and *cfg* files.

For the steps $n > 0$, UPMaBoSS runs MaBoSS according the *bnd* file and a new *cfg* file (named `cfg_file_step_n.cfg`, where `cfg_file` is the name of the initial *cfg* file). For that, an updated probability distribution is constructed from the final probability distribution of the step $n - 1$ in the following way:

1. Set to 0 each probability whose state has the death node active.
2. Double each probability whose state has the division node active.
3. Define the population ratio of step $n - 1$ as the sum of all probabilities (after applying 1 and 2).
4. Set division node to inactive for every state in the probability distribution.
5. Divide all probabilities by the population ratio.

The initial *cfg* file is copied to a new file `cfg_file_step_n.cfg`. The updated probability distribution (above) is injected in this new file as the new initial condition. The values of every external variable in the upp files are computed according to this updated probability distribution. These new values of external variables are injected in the new *cfg* file `cfg_file_step_n.cfg`.

#### 1.3 Description of output files

For the executable version of UPMaBoSS (in perl), cfg files for each MaBoSS run are generated. In addition, two files are provided: `Model_PopR.csv` and `Model_PopProbTraj.csv`.
`Model_PopR.csv` contains the population ratio at each time step.
`Model_PopProbTraj.csv` contains network state probabilities at each time step.

#### 1.4 Choice of time step length

The `max_time` value in the *cfg* file is considered as the "time step", because UPMaBoSS updates the population model after a MaBoSS run of this length. Therefore, this value needs to be carefully chosen:

- The time step should be smaller than the first transient effect.
- The time step should be large enough so that this transient effect had already been initiated.

Therefore, it is strongly suggested to first produce a single run of MaBoSS, using the *bnd* file and a modified *cfg* file that has a much longer `max_time`, in order to estimate these lower and higher limits of the "time step" described above. Then, time step sensitivity within its lower and higher limits should be studied (see the example below).

#### 1.5 Parameters to be set for a simulation

##### Parameters that have a biological interpretation

- Activation/inhibition rates: numerical values put in the formulas for `rate_up` and `rate_down` in the *bnd* file. They represent the speed of activation/inhibition of the associated node. A possible interpretation of this value is 1/time of transition. The suggested default value is 1.0. They are related to the unit of time used for the length of time steps defined below, *eg* if the unit of time steps is [hours], the rates are expressed in unit [1/hours].
- Time length of simulation. This value is set by the number of steps given in the *upp* file. The length of the simulation corresponds to this number of steps multiplied by the length of time steps (see below) defined as the length of MaBoSS simulation (`max_time` in the *cfg* file). As default, when activation/inhibition rates are close to 1, we suggest to take 10 steps for a time step length of 10.
- Initial conditions. The initial condition is set as a probability distribution over the set of network states. As default, we suggest to take a random probability distribution, where all network states have an equal probability. When modeling a specific pathway that represents *in vitro* experimental data, all node could be set at zero except the one that activates the pathway.
- Update formula for external variables in receptors. This is given by the equation, in the *upp* file, that describes the update of an external variable in term of state probabilities. Any formula using $+, -, *, /, \log, \exp$ can be used. For an external variable (eg `$outerL`) describing an activation rate of a receptor by a ligand (L), we suggest to use `$outerL u= p[(L) = 1]`.

- Time tick. This value is set in the  file (`time_tick`). It represents the length of time windows on which probabilities are estimated. Although it is preferable to use small values, a too small value will produce non-continuous probability trajectories. This value shouldn't be larger that the time window on which time dependant experimental values could be measures. The suggested default value is 0.1.

**Modeling parameters**

- Number of MaBoSS trajectories. This parameter, set as `sample_count` in the  file, controls the quality of simulation. It is recommended to increase this value if the results depend too much on the seed of the random generator. The suggested default value is 10000
- Seed of random generator. This parameter, set as `seed_pseudorandom` in the *cfg file*, allows to test the stability of the results by changing its value. There is no suggested default value.
- Length of time steps between each update. This is set in the *cfg* file by the `max_time` value, see the discussion above. The unit of time is the inverse of the unit of activation/inhibition rates. The suggested default value is 10.

### 2 Application: population modeling of cell fate model

#### 2.1 Model parameters

The population version of TNF induced cell-fate is constructed in the following way:

1. We used a published model, translated in MaBoSS language (available in MaBoSS webpage: https://maboss.curie.fr/).
2. We added a node "Division", induced by "Survival" node, that has an activation rate of 1/12 (half a day).
3. We added a node "Death", induced by "Apoptosis" or "NonACD" nodes, that has an activation rate of 1 (one hour).
4. We fixed the time step to 1 hour, because the transient activation of death has its first maximum at 1 hour.
5. For implementing TNF production by NF$\kappa$B, we added an external variable $TNF_induc. This variable is updated according to the probability of having NF$\kappa$B active and Death inactive.
6. We launched UPMaBoSS.pl with 48 steps, upon different conditions (see below).

.

We consider the following conditions:

- Transient TNF (initial state with TNF active, TNF degradation rate of 1/6).
- Transient TNF with no cell-cell interaction (no induction of TNF through NF$\kappa$B).
- No TNF (TNF inactive initially).
- Permanent TNF ($TNF_induc at 20 in the initial *cfg* file), starting from the asymptotic state from the previous UpPMaBoSS run (transient TNF or no TNF).

#### 2.2 Time step sensitivity

A Jupyter notebook (`TimeStepDependency.ipynb`) describes the effect of changing the max_time for two conditions when TNF is ON (TNF) and when TNF is OFF (NoTNF). The different population ratios are compared for the values: 1 (the value used for our example), 0.5, 0.33, and 0.25. This example is available within the CoLoMoTo Docker image at https://github.com/sysbio-curie/UPMaBoSS-docker.
